# Predicting functionally important breast cancer SNPs using pleiotropy, conservation, and protein structure

**DOI:** 10.1101/2024.01.01.573831

**Authors:** Meredith A. Carpenter, Alan C. Cheng

## Abstract

**Motivation:** With over 24,000 SNPs associated with breast cancer in ClinVar, there is a need to prioritize the subset most likely to be causally linked to diagnostic, prognostic, and other clinical outcomes of disease. Building off currently known breast cancer oncogenes and SNPs, we identify the subset of SNPs with pleiotropic effects, with the goal of identifying mutations functionally relevant to disease progression. We further use sequence and structure analysis to prioritize missense mutations most likely to impact protein function.

**Results:** From the known breast cancer SNPs, we identified co-associated mutations located at evolutionarily conserved positions and contributing significant protein stability as potential focal points for disease biomarkers, protein function studies, and therapeutic intervention. To identify regions likely integral to protein function, we plotted genomic intervals where multiple disease density peaks overlap. Of the breast cancer SNPs, 1,714 were co-associated in-frame mutations, of which 930 occurred at conserved residue positions (Shannon Entropy <1.0) and 833 were also missense mutations. Building structure-based models of the 277 SNPs with available protein structure resulted in identification of 133 SNPs that are calculated to affect protein thermostability by >100-fold (>3 kcal/mol). The workflow we built can be applied to other diseases to help identify functional mutations.

**Availability:** Python code for the integrated analysis workflow available at http://github.com/mcarpenter-brandeis/brc-pleiotropy and detailed data tables available in Supplemental Information.

## 1 Introduction

Breast cancer is a widespread, complex disease with high variability in tumor pathology as well as high genomic and epigenomic diversity in affected individuals (Polyak, 2011). Genetic contributions to carcinoma progression are most often the result of germline mutations in genes involved in pathways integral to genomic integrity and, as a result, cellular functioning (Walsh and King, 2007). Mutations in key tumor suppressor genes and oncogenes underlie both somatic and heritable familial breast cancers, with the latter constituting 15-20% of breast cancer occurrences (Wendt and Margolin, 2019).

No singular gene mutation is known to guarantee a breast cancer phenotype. Instead, numerous distinct mutations can lead to increased relative risk and polygenic risk of disease occurrence and recurrence, and differing responses to interventions (Keen and Davidson, 2003; Cavalieri *et al*., 1997). To date, dozens of breast cancer susceptibility genes have been identified. The most extensively investigated are BRCA1 and BRCA2, with over 1000 different mutations characterized on these two genes (Antoniou and Easton, 2006). Though mutations on these two genes are estimated to account for up to 80% of the heritable component of familial breast cancers when present (Greene, 1997), BRCA1 and BRCA2 mutations only account for ~5% of total familial breast cancers.

Other moderate risk susceptibility genes include CHEK2, which was identified in a non-BRCA family (Wendt and Margolin, 2019), and ATM, PALB2, and RECQL, which were characterized by familial genomics and cohort studies (Cybulski *et al*., 2015). Low-risk breast cancer variants have primarily been identified through GWAS studies (Wendt and Margolin, 2019). Subpopulation and twin studies, including of European, Icelandic, Finnish, Ashkenazi Jewish and East Asian populations have been useful in identifying disease-relevant loci and for calculating population attributable risks of deleterious mutations (Hemminki and Czene, 2002). Currently, over 2500 RefSeq transcripts have NCBI ClinVar entries annotated with breast cancer or breast cancer-associated conditions, resolving to approximately 270 unique, non-overlapping genes (Landrum *et al*., 2018). The subset of genes that increase heritable risk for breast cancer and used in this study are in Table S1.

Breast cancer associated genes generally are involved in genomic stability, DNA repair, cell cycle regulation, and apoptosis. Many are proto-oncogenes, oncogenes, and tumor suppressor genes, and are associated with carcinomas, malignancies, and cancer syndromes. GWAS studies have identified pleiotropic associations between various cancers, including between breast cancer and lung, prostate, and ovarian cancers (Fehringer et al., 2016). In addition, breast cancer genes have been found to be associated with multiple cancer predisposition syndromes (i.e., Li-Fraumeni, PTEN hamartoma tumor, Cowden, and Peutz-Jeghers syndromes) and additional cancers (pancreatic, colon, endometrial, lung, gastric, bladder, lymphoma, leukemia, neuroblastoma, Ewing’s sarcoma, glioblastoma, glioma, medulloblastoma, osteosarcoma, and melanoma). Table S1 details disease co-associations for breast cancer genes.

As genomic databases grow through public access screening and directed mutational studies, methods for identifying genes and variants that cause disease are needed. These curated pathogenic variants can be useful for diagnostics, and, potentially, to guide therapeutic intervention. Here, we leverage non-oncogenic disease annotations, sequence conservation analysis, and structure-based modeling to identify likely pathogenic variants with a high degree of corroborating evidence.

We first identify breast cancer susceptibility genes that contribute to non-oncogenic diseases, including blood, kidney, cephalic, neurologic, musculoskeletal, and additional systemic disorders listed in Table S2. A prime example of a non-oncogenic condition co-associated with numerous breast cancer variants is Fanconi anemia (FA). FA is a recessive syndrome characterized by chromosomal instability and hypersensitivity to agents that produce DNA interstrand crosslinks (ICLs) such as chemotherapy drugs. FA proteins cooperate in a pathway leading to recognition and repair of damaged DNA, and they interact with other proteins integral for chromosome stability (Taniguchi and D’Andrea, 2006). Mutations in the FA genes thus have pleiotropic effects and result in both FA and inducement of considerable breast cancer risks like those observed with BRCA1 or BRCA2 mutations (Howlett *et al*., 2002).

This study uses additional variants similarly identified as having pleiotropic linkage. Candidate mutations are further evaluated with respect to their genomic distribution patterns, evolutionary conservation at specific residue positions, and impact on protein stability, with the goal of identifying the subset of breast cancer SNPs most likely to result in protein dysfunction and, therefore, disease. We make available our code to enable further analysis as well as application to non-breast cancer diseases (link provided in Abstract).

## 2 Methods

## 2.1 Identifying candidate SNPs

Breast cancer associated genes were queried in the ClinVar database to identify all SNPs annotated on that gene, breast cancer associated or otherwise (See Table S1 for queried genes; data sets obtained on June 29, 2021). Secondary diseases were extracted from this SNP dataset as the non-cancer conditions annotated on breast cancer annotated SNPs— ClinVar condition(s): breast cancer + secondary condition. The full gene set considered in this study was derived from the pleiotropic SNP dataset as any gene containing co-associated SNP(s).

When there are multiple RefSeq transcripts associated with a variant, ClinVar reports all associated transcripts. Variants associated with more than two transcripts were excluded from consideration due to lack of resolution to a specific gene. Most variants associated with two transcripts had the query gene as the first gene listed in ClinVar and were thus assigned to that gene. The remaining variants associated with two transcripts had the query gene listed second. These mutations were evaluated and were able to be confidently assigned to the query gene. Specifically, C11orf65|ATM SNPs (rs1565520641, rs1555117132, rs563137460, rs786203311, rs587781344, rs786202826) were assigned to ATM as each mutation had matching OMIM entries on ATM; AOPEP|FANCC SNP rs1588070592 was assigned to FANCC because the mutations aligned with FANCC and not AOPEP. Only variants with a dbSNP identifier were used for co-association analysis and calculations (entropy and protein stability). Genes used in the study are in Table S1.

## 2.2 Identifying regions of SNP co-occurrence

Density plots were generated for each of the genes of interest with the GrCh38 locations from ClinVar plotted as the x-axis values. Mutations that span a location region interval were placed at the mean chromosome location value 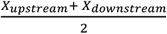 where *XX*_*upstream*_ and *XX*_*downstream*_ define the mutation region.

Plots contain all SNPs on the genes of interest including individually listed diseases (breast cancer or secondary disease) and co-associated diseases (breast cancer and secondary disease). SNPs annotated with multiple disease conditions were counted independently for each disease. “Other Cancer” refers to any SNP with a specific cancer excluding breast cancer and hereditary cancer-predisposing syndrome. For any gene containing at least two qualifying entries, subset plots were generated and colored by condition. Density plots were generated using the Seaborn distplot function without histogram (hist=False), with kernel density estimation (kde=True), and with a legend labeled and color coded by condition. Plots were generated using Python 3.9.0 and Seaborn (Waksom, 2021). Conditions with only one SNP on the gene were not plotted since kernel density estimations (KDEs) cannot be produced for a single data point.

### 2.3 Heatmap of SNP abundance and distribution

A heatmap was generated to visualize the general distribution and abundance of co-associated SNPs across the genes of interest using the ClinVar database derived SNP information. The number of SNPs for each disease was calculated for each of the genes containing at least one co-associated SNP (Table S3). The column “Breast Cancer” represents all SNPs that contain a breast related disease including, but not limited to, breast cancer, breast carcinoma, familial breast cancer and hereditary breast cancer predisposition. The second column, “Other Cancer” represents SNPs with a second, specific cancer listed in addition to a breast related disease. Hereditary cancer-predisposing (HCP) syndrome was not considered a separate, specific cancer so SNPs listed with breast cancer and HCP were not counted in this second column. The remaining columns represent SNP counts of co-associated variants associated with breast cancer and the disease column label. Plots were generated using Python 3.9.0 and the Seaborn library.

### 2.4 Generating SNP variant sequences and Shannon Entropy conservation calculations

Sequences were aligned on a single isoform (most often isoform 1) catalogued in UniProt (The UniProt Consortium, 2021). Amino acid variants were derived from the ClinVar entries ‘Protein Change’ column. ClinVar protein changes are listed as: Isoform AA _ Residue Number _ SNP mutation AA. For each SNP, the listed pre-mutational amino acid was matched to the canonical isoform at the given residue position before the sequence was mutated to the SNP amino acid. In a few cases, SNP information in the dataframe had to be adjusted to coincide with the annotations in ClinVar (noted in Python code). In-frame SNP mutations, including missense and nonsense (*) were used, while frameshifts (fs), insertions (ins) and deletions (del) mutations of any length were excluded. BLAST (Altschul *et al*., 1997) was used to identify homologs (NCBI NR, up to top 1000 sequences, E < 1.0 x 10^-40^) (Pearson, 2013) for each gene. MSAs of the query with homologous sequences were used to generate counts of amino acids at each identified SNP residue. Shannon entropy (SE) was calculated as:

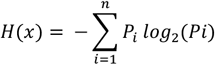

Homologs with gaps at the evaluated residue were excluded from the SE calculation. SE and alignment count for each evaluated SNP are reported in Table S6. Shannon entropy conservation values presented for the eight examples in Fig. 4 exclude homologous sequences where the residue aligned is a gap; the percent excluded sequences for the eight SNPs are (A) ATM 51%, (B) BLM 4%, (C) BRCA1 33%, (D) BRCA2 64%, (E) FANCC 25%, (F) HRAS 4%, (G) mTOR 50%, (H) PALB2 26%.

### 2.5 Rosetta calculation: Effect of SNPs on protein stability

We selected PDB structures for proteins of interest based on completeness, resolution, and sequence identity (>75%) (Berman *et al*, 2000). For our analysis, we removed waters, ligands, DNA, and other non-protein components. Missing loops and gaps in protein structures were homology modeled using SWISS-MODEL (Waterhouse *et al*, 2018) (Table S4). Of the 833 conserved, missense SNPs scored for SE, 277 had positions resolve to the protein structure and wild-type amino acid match the model residue at the location. The structures were mutated in PyRosetta, according to their SNPs, and then minimized using FastRelax with the ref2015 potential. Relaxed PDB files for all genes are provided in Supplemental materials. A SNP’s effect on protein stability was calculated as ∆*E =E*_*SNP model*_*−E*_*WT model*_ where *E*_*SNP model*_ = pose energy score calculated after SNP mutation and *E*_*WT model*_ = pose energy score of the unmutated protein. SNPs with a change in energy (∆E) > 3.0 kcal/mol were predicted to decrease structural stability. Models were visualized in PyMol (Schrödinger Inc.; DeLano, 2020).

## 3 Results

We looked at ~33,800 SNPs currently listed in ClinVar as associated with breast cancer (~24,700 with associated dbSNP IDs). In the broadest analysis of the data set, density plots were generated with SNPs associated with either breast cancer, at least one secondary disease of interest, or both. Pleiotropic, co-associated SNPs used in the remainder of the study (2,726 SNPs associated with one of 45 genes) were specifically those SNPs that have both breast cancer and a non-cancerous secondary condition of interest listed in ClinVar.

Excluding insertions, deletions, and frameshifts resulted in 1714 co-associated, in-frame mutations (missense and nonsense), of which 930 SNPs (34 genes) had a Shannon entropy less than 1.0. All conserved, nonsense mutations evaluated were indicated in the literature to be pathogenic because of premature translational stop that likely results in a loss of function through truncation or nonsense-mediated mRNA decay. Missense mutations have a more variable effect depending on the location, function and relative conservation of the amino acid being mutated and were thus further evaluated. Of the 930 conserved, in-frame SNPs, the 833 missense mutations were considered for mapping to structure and 277 SNPs across 27 genes were mapped to a relevant PDB protein structure. We used Rosetta to predict the degree of protein destabilization resulting from each SNP. Of the 277 SNPs, 133 mutations were predicted to have a significant effect on protein structure destabilization (∆E > 3.0 kcal/mol). This approach integrates analyses on co-associated diseases, evolutionary conservation, and protein stability highlights potentially significant mutations by narrowing down the initial dataset to less than 1% of SNPs.

### 3.1 Pleiotropic gene regions and SNPs

To better understand the chromosomal distribution of SNPs from breast cancer and secondary diseases, we generated SNP density plots for all 45 genes considered in this study (Supplemental materials). Fourteen genes had only a single breast cancer annotated SNP so density curves could not be plotted for the primary disease. Three genes had only one SNP annotated with a secondary (non-breast cancer) disease. Both breast cancer and at least one secondary disease curve could be calculated for the remaining 28 genes. Peak regions, specific SNP locations and rsID numbers for co-associated SNPs on all evaluated genes are summarized in Table S5. These density plots highlight regions of the genes that have high densities of co-associated SNPs and may identify regions of the gene that are important for protein function.

Example density plots for ATM, HRAS, PIK3CA and TSC1 are shown in Fig. 1 and can offer a visual perspective of SNP density distributions tied to multiple disease conditions. SNPs on ATM annotated with breast cancer and ataxia, a condition where one loses control of bodily movements, have a highly similar distribution, suggesting that SNPs in these regions contribute to both disease pathologies. Low-tailed curves, such as for microcephaly on the same plot, suggest SNPs in only a narrow region are associated with both diseases. Conversely, highly tailed curves suggest SNPs in dispersed regions are associated with the condition, such as immunodeficiency and seizure also on ATM. In the case of HRAS, the breast cancer associated SNPs are tightly clustered, while the Costello syndrome and Noonan syndrome associated SNPs are more widespread across the gene. PIK3CA has three density peaks, two of which are shared by all conditions annotated on the gene, but one of which (the central peak) is unique to breast cancer SNPs. Overall, the distribution of SNPs on the genes are frequently clustered together but can differ in the tightness of overlap between the specific disease curves. BRIP1 for example has nearly identical distributions of breast cancer and FA SNPs, suggesting that most, if not all SNPs on this gene are co-associated FA-breast cancer SNPs. TSC1 provides a counterexample where all the density curves are offset and separated, indicating much less overlap and co-association of SNPs on that gene.

**Fig. 1.**
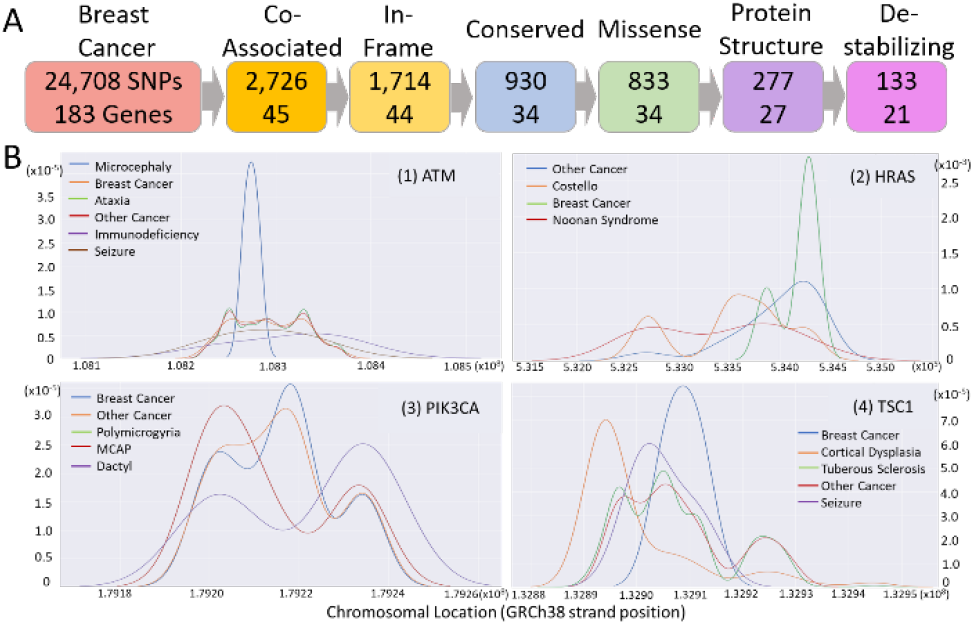
Workflow and selected SNP distributions. (A) Workflow depiction with number of SNPs and genes identified/analyzed at each step. Conserved SNPs have Shannon Entropy <1.0 and predicted destabilizing SNPs have calculated ∆E stability score >3.0 kcal/mol. (B) Distribution plots with kernel density estimation (kde) based on GrCh38 chromosome locations for ClinVar SNPs associated with identified conditions for selected genes, (1) ATM, (2) HRAS, (3) PIK3CA, and (4) TSC1. Conditions with 1 or fewer SNPs on the genes are not plotted. Plots generated using Seaborn.

While the density distribution plots allow identification of regions contributing to multiple diseases, we were also interested in individual SNPs that are implicated in both breast cancer and a secondary condition. Fig. 2 provides a heatmap view of these SNPs. The distribution trends parallel the density plots but with lower frequencies due to the additional requirement of co-association. In Fig. 2, we see that BRIP1, ATM, BRCA2, PALB2, BRCA1, PIK3CA, CDH1 and HRAS are abundant in both breast cancer and multiple cancer (breast + secondary cancer) SNPs. RAD51C, NBN, FANCC, BLM, FANCM, MRE11 and RAD50 also have many breast cancer SNPs, however, are not as generally oncogenic in this analysis as they have few SNPs co-associated with multiple cancers. Consistent with the density plots, several genes have only a single SNP associated with both breast cancer and one or more secondary conditions; these genes fall within the lower half of the heatmap.

**Fig. 2.**
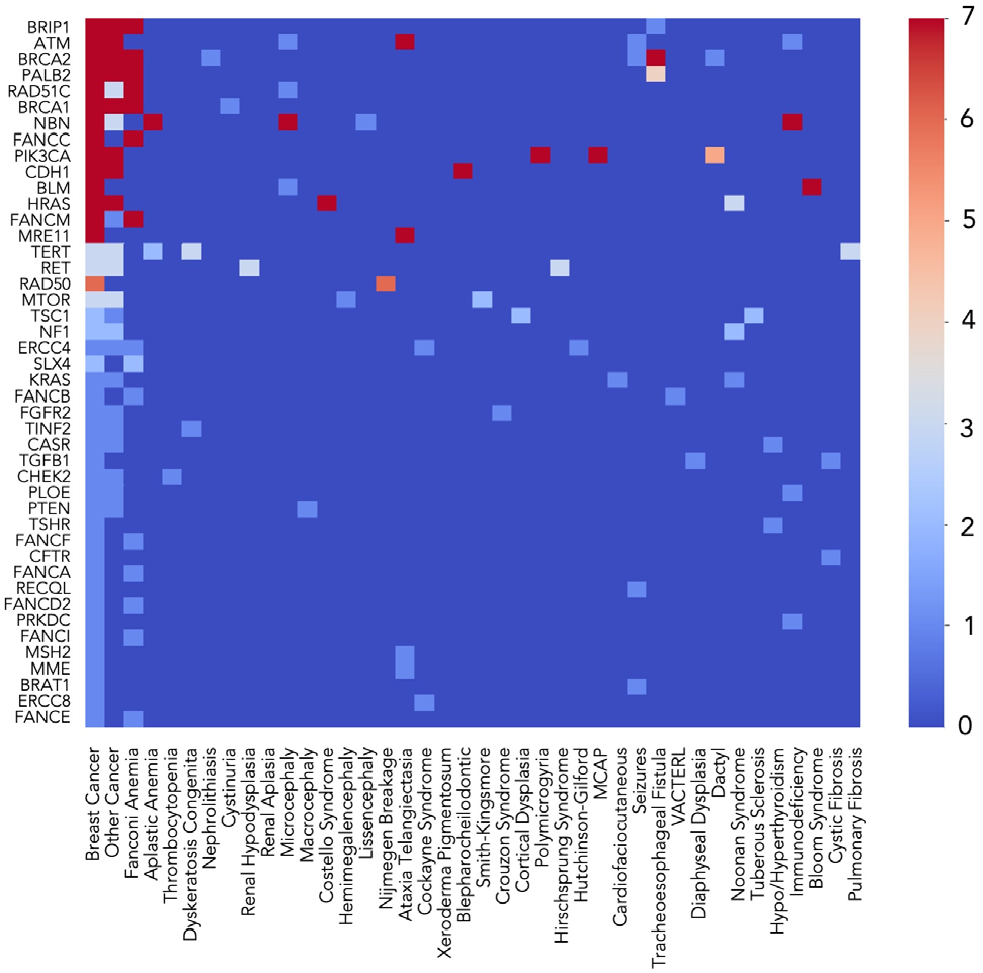
Heatmap of co-associated SNP counts. All SNPs in our count database are classified under Breast Cancer. “Other Cancer” includes co-associated SNPs with at least one non-breast associated cancer additionally listed. All other SNP counts indicate any SNP annotated with the disease listed in the column. Genes are ordered from highest (top) to lowest (bottom) cumulative number of SNPs. MCAP = Megalencephaly-capillary malformation; VACTERL = vertebral defects, anal atresia, cardiac defects, tracheo-esophageal fistula, renal anomalies, and limb abnormalities.

Aside from breast cancer, Fanconi anemia is the most abundant non-cancerous secondary disease both with respect to the number of SNPs and number of associated genes. SNPs associated with any Fanconi anemia complement group were identified on BRIP1, BRCA2, PALB2, RAD51C, BRCA1, ERCC4, SLX4, and the FANC genes (FANCA, FANCB, FANCC, FANCD2, FANCE, FANCF, FANCI and FANCM). All other secondary conditions are annotated on four or fewer genes, usually one to two genes. Four conditions are noted on four different genes: Microcephaly (ATM, RAD51C, NBN, BLM), Ataxia telangiectasia (ATM, MRE11, MSH2, MME), seizures (ATM, BRCA2, RECQL, BRAT1), and immunodeficiency (ATM, NBN, POLE, PRKDC). Two are noted on three genes: Transesophageal fistula (BRIP1, BRCA2, PALB2) and Noonan syndrome (HRAS, KRAS, NF1).

### 3.3 Residue conservation and protein thermostability stability calculations

SNP positions with Shannon entropy ≤ 1.0 were considered as conserved (further details in Methods). The results of our analysis are summarized in Table 1. Alignments and alignment description tables are provided as a zip file in Supplemental materials. Expanded Shannon Entropy (SE) scoring tables are available in Table S6.

**Table 1:**
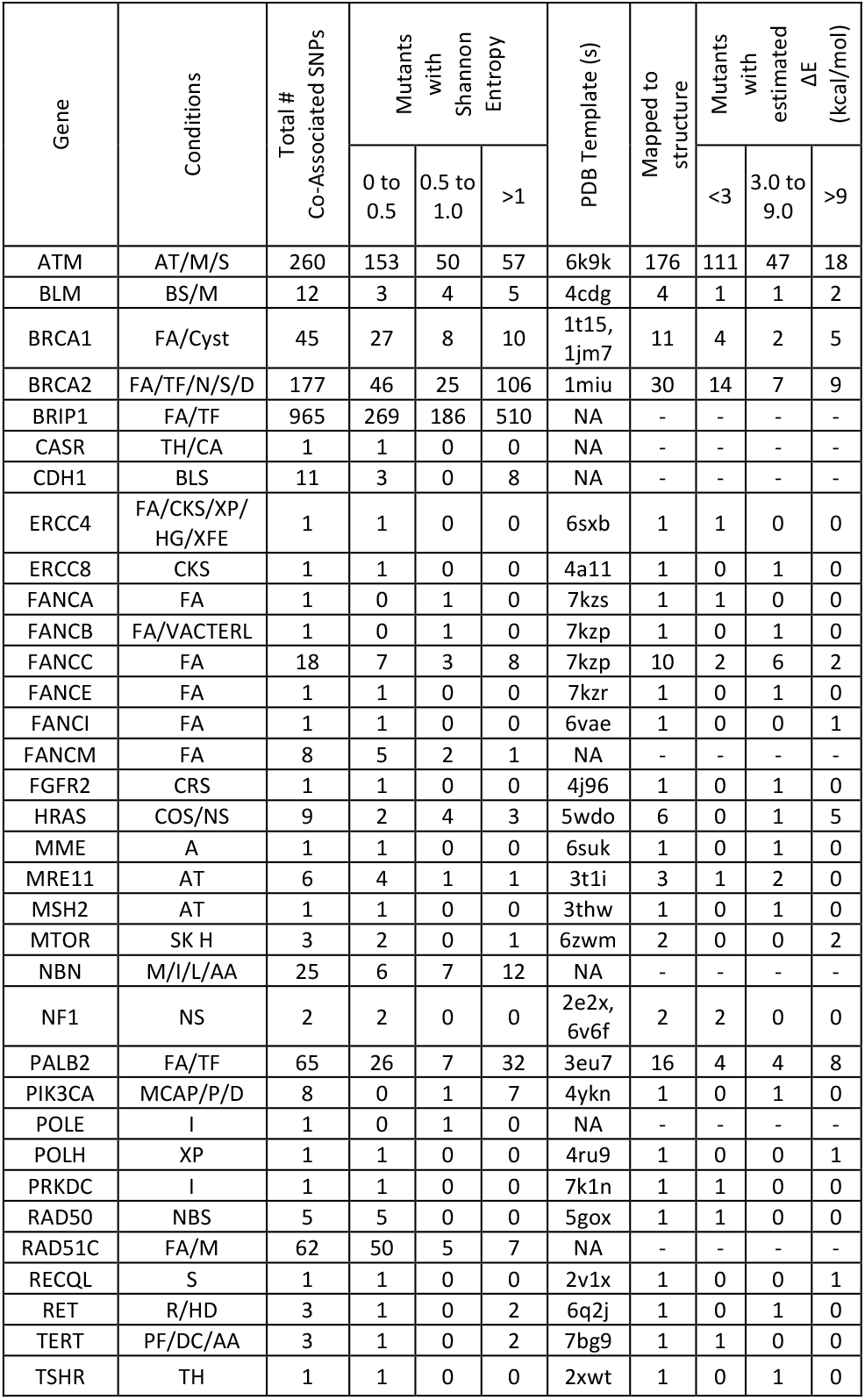
Shannon Entropy and protein modeling results. In total 930 of 1714 in-frame variants were located at conserved positions (SE < 1.0) on 34 of 45 genes. Genes with no conserved SNPs not shown. Secondary conditions associated with genes are noted alongside counts of co-associated SNPs and conserved SNP by SE. PDB templates used for homology modeling are reported if there is at least one mapped SNP. Condition abbreviations: A = Ataxia, AT = Ataxia Telangiectasia, M = Microcephaly, S = Seizures, BS = Bloom Syndrome, FA = Fanconi Anemia, Cyst = cystinuria, TF = Tracheoesophageal fistula, N = Nephrolithiasis, D = dactyly, TH = hypo/hyperthyroidism, CA = Hypo/hypercalcemia, BLS = Blepharocheilodontic syndrome, CKS = Cockayne Syndrome, XP = Xeroderma Pigmentosum, HG = Hutchinson-Gilford, XFE = XFE progeroid syndrome, CRS = Crouzon Syndrome, COS = Costello Syndrome, NS = Noonan Syndrome, SK = Smith-Kingsmore Syndrome, H = Hemimegalencephaly, I = Immunodeficiency, L = Lissencephaly, AA = Aplastic anemia, P = Polymicrogyria, NBS = Nijmegen Breakage Syndrome, R = Renal hypodysplasia/aplasia, HD = Hirschsprung disease, DD = Diaphyseal dysplasia, CF = Cystic fibrosis, PF = Pulmonary fibrosis, DC = Dyskeratosis congenita.

Overall, 11 genes (BRAT1, CHEK2, CFTR, FANCD2, FANCF, KRAS, PTEN, SLX4, TGFB1, TINF2, TSC1) did not have any conserved co-associated SNP residue positions, six genes had fewer than 50% of SNP positions conserved (BRCA2, BRIP1, CDH1, PIK3CA, RET, TERT), and the remaining 28 genes had most evaluated SNP positions conserved (SE <=1.0). A mean of 517 sequences were utilized to calculate conservation score at each residue, with only five residue positions scored using an alignment with less than 100 sequences. 115 residues across 15 genes had a calculated SE = 0.0 with greater than 235 sequences aligned, 41 with greater than 500 sequences aligned. In several cases, positions with SE =0.0 have multiple identified SNPs associated with different amino acid substitutions. For example, rs876659558 (E2039V) and rs864622251 (E2039K) on ATM, rs768555161 (R777C/G) and rs747568830 (R777P/H), as well as rs587780249 (D1138N), rs1555572620 (D1138Y/H) and rs1057518847 (D1138G) on BRIP1. The high conservation of the residue potentially explains why multiple amino acid substitutions at these sites are associated with disease states.

Over 40 pleiotropic SNPs occur in very highly conserved (SE <= 0.25) residue positions on BRIP1 (224 SNPs), ATM (110 SNPs) and RAD51C (42 SNPs). Including SNPs with SE <= 0.50, BRCA2 (46 SNPs), BRCA1 (27 SNPs) and PALB2 (26 SNPs) also have an abundance of conserved SNPs. Of the genes with a single co-associated SNP score, 80% (12 of 15) had a highly conserved SE (<=0.50). These include CASR rs1801725, ERCC4 rs121913049, ERCC8 rs61754098, FANCE rs145068586, FANCI rs202066338, FGFR2 rs1057519045, MME rs138218277, MSH2 rs1573578602, POLH rs2307456, PRKDC rs8178208, RECQL rs150306543, and TSHR rs142063461. Particularly in the genes with high counts of conserved SNPs, regional conservation scoring patterns are noticeable, with numerous adjacent or nearby SNPs scoring similarly. For example, BRIP1 has five SNPs with SE = 0.0, 15 SNPs with SE = 0.01 and two SNPs with SE = 0.02 between residues 802-865. Regions abundant with highly conserved SNPs are potentially functional important protein domains within which any mutation can disrupt protein activity, binding interaction, or destabilize the protein itself. To evaluate the structural importance of these conserved residues, we modeled the mutations on known protein structures and calculated the change in energy (∆E) resulting from the amino acid substitutions.

### 3.4 Mutational effects on protein stability

Of the 833 highly conserved, missense, co-associated SNPs considered in this study, we were able to model and score 33% of them using Rosetta protein design tools to assess potential destabilizing effects on the protein structure. The remaining SNPs could not be modeled due to the lack of comparable protein structure. BRIP1, NBN, POLE, and RAD51C had no relevant structures in RCSB PDB with greater than 75% sequence identity, and thus all SNPs on those genes were excluded from scoring. Conserved SNPs on CASR, CDH1 and FANCM were outside of the modeled regions on available comparable protein structures. Of the 277 modellable SNPs on the remaining 30 genes, 133 SNPs, or approximately 48%, had a change in protein stability score (∆E) > 3.0 kcal/mol, indicating a predicted decrease in protein stability because of the mutation.

Table 1 provides a summary of the predicted ∆E contributions to protein stability for SNPs on each gene. Table 2 provides an expanded table with gene, dbSNP identifier, protein change and scoring of all co-associated, missense mutations having Shannon entropy <1.0 and ∆E >3 kcal/mol. Structural models of selected SNPs with pleiotropic effect, strong sequence conservation (Shannon entropy <1.0), and calculated to be highly protein destabilizing (∆E>9 kcal/mol) are provided in Fig. 4. Stability scores for all modeled SNPs are provided in Table S6.

**Table 2:**
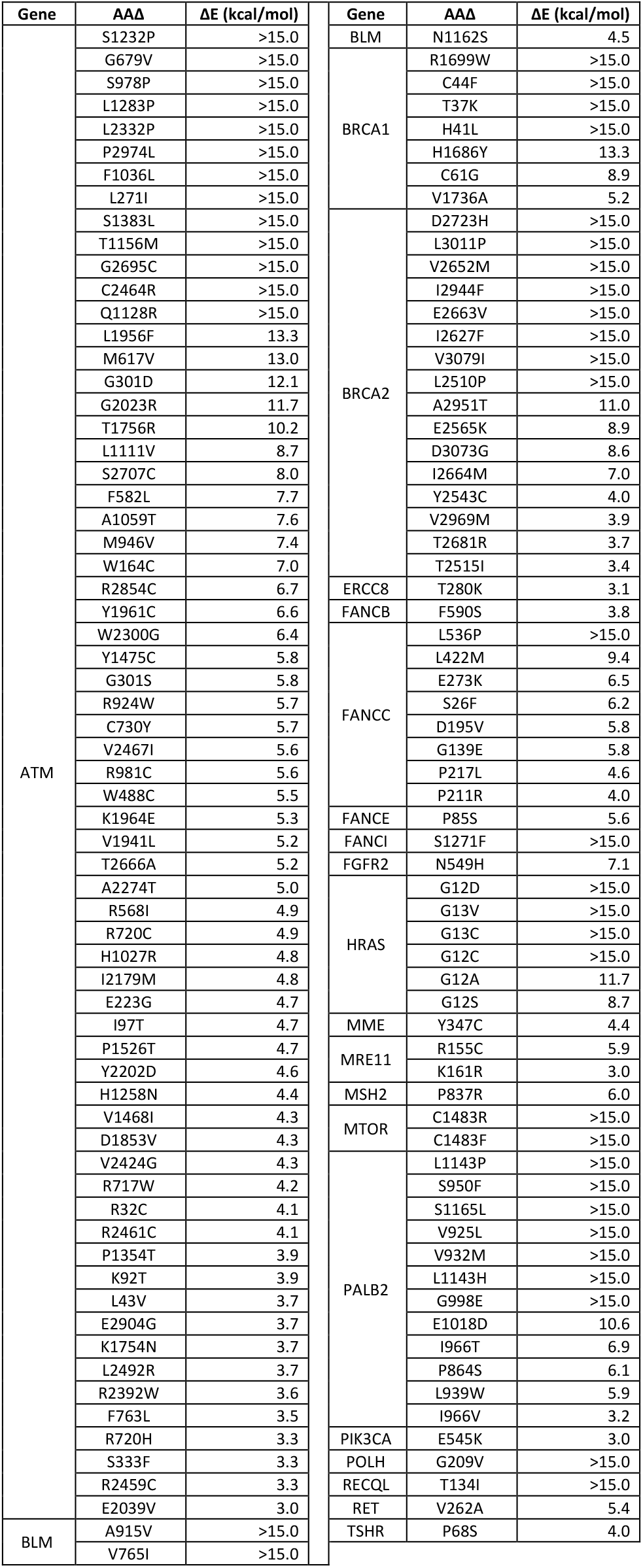
Structure-based predictions of destabilizing conserved, co-associated, missense mutations. Shown here are the 133 co-associated, missense SNPs with SE < 1.0 and computed ∆E > 3.0 kcal/mol. See Table S6 for dbSNP rsIDs with the associated protein change (ClinVar). Rosetta was used to calculate ∆*E = E*_*SNP model*_*−E*_*WT model*_ where *E*_*SNP model*_ is the energy after SNP mutation and *E*_*WT model*_ is for the unmutated protein.

Twenty-seven genes had at least one SNP identified by our approach, and five genes had ten or more SNPs modeled. Greater than half of the mutations evaluated on FANCC (80%), PALB2 (75%), BRCA1 (64%) and BRCA2 (53%) were predicted to be destabilizing as defined by a ∆E > 3.0 kcal/mol. Conversely, most scored mutations on ATM (111 out of 176) had ∆E < 3.0 kcal/mol, suggesting that while there is an abundance of noted mutations on ATM, there is a high variability in their potential pathogenicity.

All together our analysis seeks to identify breast cancer SNPs with the largest effect on disease based on genetic data, sequence analysis, and protein modeling. We specifically look at multiple functional effects (pleiotropy), residue conservation among homologs, and calculated protein stability leveraging available protein crystal structures. The full analysis highlights 133 SNPs on 21 proteins that occur in highly conserved residue positions (Shannon entropy <1.0) and result in a significant calculated change in protein stability (∆E >3 kcal/mol). To better understand our findings, we describe eight examples of SNPs identified by the complete workflow, shown in Fig. 3, and the full list of 133 is available in Table S6.

**Fig. 3.**
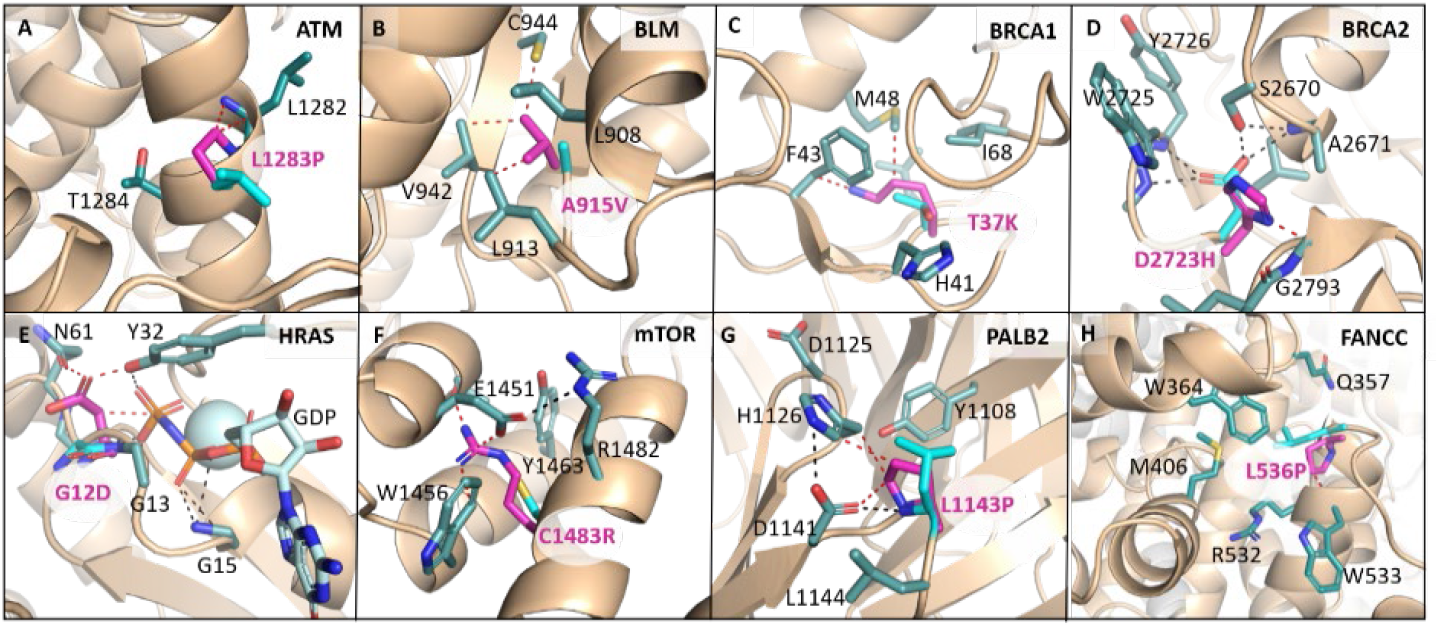
Structural models of selected SNPs. SNPs identified based on pleiotropic effect, sequence conservation, computed change in protein stability, as well as crystal structure availability. The eight examples shown here all have calculated stability ∆E>9.0 kcal/mol and alignment position Shannon Entropy<1.0. (A) ATM rs730881389 (L1283P) modeled based on PDB:6k9k, (B) BLM rs775026151 (A915V) on PDB:4cdg, (C) BRCA1 rs80356880 (T37K) on PDB:1jm7, (D) BRCA2 rs41293511 (D2723H) on PDB:1miu, (E) HRAS rs104894230 (G12D) on PDB:5wdo, (F) mTOR rs1057519914 (C1483R) on PDB:6zwm, (G) PALB2 rs62625284 (L1143P) on PDB:3eu7, (H) FANCC rs587779903 (L536P) on PDB:7kzp. Wild-type variant is shown in wheat and teal/cyan stick, and SNP variant mutation is modeled in magenta. Black dashed lines indicate favorable hydrogen bonds in the wild-type variant, and red dashed lines indicate steric clashes in the SNP variant.

Our first example, shown in Fig 3A, occurs in ATM and involves a SNP, rs730881389 (CTA>CCA), that results in a L1283P mutation in the N-terminal solenoid domain. ATM is a regulator of DNA damage response (Xiao *et al*., 2019) and L1283 is 100% conserved, excluding gaps, in all 421 homologs. The mutation is annotated as likely pathogenic in ClinVar and has been observed in at least three individuals with Ataxia-telangiectasia as well as in individuals affected with breast and ovarian cancer (Mitui *et al*., 2005).

Our second example, shown in Fig 3B, occurs in BLM, which encodes a DNA helicase involved in homologous recombination. The SNP, rs775026151, results in an A915V change in the helicase C-terminal domain (Swan et al., 2014). A915 is 100% conserved, excluding gaps, in the 963 homologs found and the variant is linked to Bloom syndrome, hereditary breast and ovarian cancer syndrome, and, potentially, Crohn’s disease (Kaneyasu *et al*., 2020). While Val is only slightly larger than Ala, structure-based modeling of A915V highlights that Ala is a protein core residue, and mutation to Val introduces several steric clashes indicated by red dashes in Fig 3B.

Our third example, shown in Fig 3C, occurs in BRCA1, which encodes an E3-ubiquitin ligase that regulates cellular responses to DNA damage. The SNP, rs80356880, covers T37K and T37R mutations in the BRCA1 RING finger domain, and the mutation of a small, slightly polar Thr to a larger Lys or Arg results in steric clashes in the core of the RING domain, as shown in Fig 4c. The mutation was modeled in a well-resolved NMR structure (PDB=1jm7), and the modeling was confirmed in recent cryo-EM structures (PDB=7jzv and 7lyb; data not shown). Nearly all homologs, 99.6% of the 686 found (excluding gaps), have a conserved Thr at the residue position. The SNP is associated with breast, ovarian, and pancreatic cancer as well as Fanconi anemia, and affects BRCA1 protein function (Starita *et al*., 2015).

Our fourth example, shown in Fig 3D, occurs in BRCA2, a tumor suppressor gene. The SNP, rs41293511 (GAT>CAT), results in a D2723H mutation in the hydrophobic core of the OB2 oligonucleotide-binding domain of BRCA2, as shown in Fig. 3D. D2723 is conserved in all 355 orthologs we identified, and D2723H as well as a related D2723N mutation have been classified as pathogenic based on studies showing the mutation affects the DNA repair function of BRCA2 (Goldgar *et al*., 2004). In addition to breast and ovarian cancers, the mutation has been linked to pancreatic and prostate cancer, medulloblastoma, glioma, Wilms tumor, and Fanconi anemia (Richards *et al*., 2015).

Our fifth example, shown in Fig 3E, occurs in HRAS, a GTPase involved in regulation of the MAPK/ERK proliferation pathway. Six SNPs occur at the highly conserved G12 and G13 positions (SE of 0.88, 86% G, and 0.27, 97% G, respectively) and the resultant protein mutations (G12A, G12D, G12C, G12S, G13C, and G13V) are calculated to cause substantial protein destabilization, with a ∆E > 8.5 kcal/mol (see Table S6). The SNP resulting in G12D (rs104894230) is calculated to have the greatest protein destabilization effect and is depicted in Fig. 3E. All six mutations have been classified by expert panels as pathogenic variants associated with a multitude of cancerous and non-cancerous indications, including neoplasm of the thyroid gland, large intestine, and thyroid gland, as well as melanoma, RASopathies, Costello syndrome, congenital myopathy with excess muscle spindles, urinary bladder cancer, and epidermal nevus (Gelb e*t al*., 2018).

Our sixth example, shown in Fig. 3F, occurs in mTOR, which is a central regulator of cell growth and cell survival processes as well as actincytoskeletal rearrangement. The SNP, rs1057519914, results in a C1483R mutation at a residue position that is highly conserved (SE = 0.04, 99.6% C), and is calculated to result in substantial protein destabilization at ∆E > 15 kcal/mol, due to the exchange of a small, slightly polar residue for a much larger, charged residue, resulting in disruption of a protein hydrophobic core. The mutation is a known pathogenic mutation that results in gain of function of the mTOR protein (VariO:0040) and is clinically associated with hemimegalencephaly, breast neoplasm, renal cell carcinoma and glioblastoma (Chang *et al*., 2016).

Our seventh example, shown in Fig. 3G, occurs in PALB2, which mediates homologous recombination by binding to and colocalizing with BRCA2 in nuclear foci, promoting stability in nuclear structures and facilitating recombination repair and checkpoint functions (Xia *et al*., 2006). The SNP, rs62625284, is associated with two protein changes, L1143P, shown in Fig. 4g, and L1143H. Leu1143 is highly conserved, with 97% of the identified homologous sequences (excluding those with gaps) having Leu at the aligned position. The L1143P mutation, shown in Fig. 3G, not only introduces steric clashes in the core of the WD40 domain of PALB2 (Oliver *et al*., 2009), but also restricts the hairpin loop in blade 6 of the PALB2-C ***β***-propeller. The protein changes are reported in ClinVar as having uncertain significance, however the L1143P mutation has been shown to reduce DNA double-stranded break-induced homologous recombination by affecting PALB2 protein interactions with RAD51C, XRCC3 and BRCA2 proteins and thereby increasing cellular sensitivity to ionizing radiation (Park *et al*., 2014).

Our final example, shown in Fig. 3H, occurs in FANCC, which is involved in apoptosis and DNA repair. The SNP, rs587779903, results in a L536P mutation at a highly conserved residue position (SE=0.01, 99.1%) which is calculated to have a destabilizing effect on the protein (∆E>15 kcal/mol) based on modeling using a cryoEM structure of FANCC (PDB 7kzp, 3.1A resolution). The substitution of L536 with a rigid Pro in FANCC is expected to disrupt the C-terminal alpha helical bundle and has been shown in one study to abolish functional activity of FANCC (Gavish *et al*., 1992). The missense mutation is classified in ClinVar as having uncertain significance (Nykamp *et al*., 2017).

We have highlighted here eight SNPs with expected significant impact on protein function. A complete list of 133 SNPs identified to affect protein function are listed in Table 2 with additional details, including rsIDs, in Tables S6.

## 4 Discussion

Clinical annotations, residue conservation, and protein stability modeling are useful tools that can be used individually or in combination in a pipeline for prioritizing the probable pathogenicity of SNPs. Identifying pleiotropic mutations or regions of disease overlap on genes can elucidate key functionally relevant regions important in multiple disease pathways. As an example, one of the most described overlapping non-cancerous conditions with breast cancer is Fanconi anemia, identified here on SNPs co-associated with breast cancer on 15 different genes. The functional association of these FA/BRC genes is well described in the Fanconi anemia (FA) complex (Walsh and King, 2007) in which the FA proteins (A, B, C, E, F, G, L, and M) activate FANCD2 via mono-ubiquitination to translocate to damage-induced nuclear foci containing BRCA1, BRCA2 and RAD51. ATM, p53, CHEK2, BRIP1, PALB2, NBS1, RAD50, MRE11 and PTEN also interact with the FA protein complex and were all identified in this analysis as containing SNPs associated with both breast cancer and non-cancerous secondary diseases. The complexes formed from these genes could explain the pleiotropic nature of the mutations, and distinct complexes can be therapeutic targets. For example, the FA E3 ligase core complex has been explored as a compelling drug target for cancer therapy, with FA pathway inactivation demonstrated to eliminate certain cancer cells without exogenous DNA damage, including cells that lack BRCA1, BRCA2 and ATM (Sharp *et al*., 2021). Focusing on pleiotropic SNPs has the potential to expand our understanding of protein complexes and pathways, and provide explanations for repeated co-occurrence of, or predisposition to, sets of conditions. However, this study and future studies are limited by the accuracy and availability of SNP annotations, as well as inconsistency of annotations and mutation identification across databases.

Identification of structurally and functionally important residues has immediate relevance to understanding protein interactions and predicting pathogenic dysfunction of proteins that can result from mutations in these key regions. Highly conserved regions are well established as indicators of residues involved in ligand binding, protein interactions, structural integrity, and functional specificity (Capra and Singh, 2007). Over half (~54%) of the SNPs evaluated in this study were located at conserved (Shannon entropy ≤1.0) and highly conserved residue positions (SE≤0.5), including 115 with SE = 0.0 as identified by 100% sequence identity of up to 1000 homologous sequences. Though conservation alone cannot indicate why the site is important, it can be used along with orthogonal analyses, such as protein structure modeling, to predict the functional relevance and potential consequences of mutations.

We used modeling to systematically and quantitatively characterize mutations that affect individual protein stability. Predicted destabilizing (∆E ≥ 3.0 kcal/mol) mutations modeled and shown in Fig. 4 tended to be sterically bulky amino acid mutations that affect protein cores or protein domain stability (such as Cys to Arg, Ala to Val, and Thr to Lys) or conformationally rigid mutations to Pro that affect both sterics and flexibility at conserved residue positions. Modeling that leverages 3D protein structure and protein stability calculations (Stein *et al*., 2019) helps in prioritizing and predicting which missense mutations are most likely to result in the loss of protein stability and the origin of disease.

Both evolutionary conservation (SE) and protein stability (∆E) were important in this pipeline, and we found little correlation (R^2^ = 0.03) between the two values across all evaluated mutations. Our approach looking at function, conservation, and protein stability is intended to identify a subset of missense SNPs that are highly likely to be functionally important. There are three main limitations to our approach here. First, we are limited by the availability of relevant protein structures. This is likely to be less of a limitation over time.

Second, we focus specifically on variants with reported pleiotropic effect, and a future study could remove this limitation albeit with expected decrease in probability of functional importance. As a proof of concept, we looked at five breast cancer specific, non-pleiotropic SNPs in PALB2 experimentally identified as destabilizing (Wiltshire *et al*., 2020; Park *et al*., 2014) were evaluated in our SE and Rosetta pipeline. The five reported SNPs, rs876658653 (L24S), rs141047069 (L35P), rs201817103 (I944N), rs863224785 (L1070P), and rs876660109 (T1030I) all had SE scores of ≤0.1 (In order listed, SE = 0.00, 0.02, 0.09, 0.04, 0.10). Three of the five SNPs (rs863224785, rs201817103, rs876660109) fell within the residues resolved in the available PALB2 protein structure and the three were calculated to be significantly destabilizing with ∆E > 8 kcal/mol. This example helps support the predictability of our pipeline.

The third limitation is that missense SNPs that affect intermolecular interactions with other proteins are largely not captured here. Highly conserved residue positions may be captured implicitly, but our structural analysis is limited by our focus on individual proteins. There are few high-resolution complex structures for the proteins of interest here. However recent advances in cryoEM structural biology are increasingly yielding atomic structures of biological complexes, and these structures can enable meaningful extension of our analysis. Advances in modeling tools such as AlphaFold (Jumper e*t al*., 2021) and RoseTTAfold (Baek *et al*. 2021) for highly accurate protein and dimer structure predictions will enable modeling of more mutations. We open-source our computational workflow (see Abstract) to enable future efforts.

Ultimately, the results are useful in prioritizing SNPs for experimental validation, such as that described for PALB2 (Wiltshire *et al*., 2020; Park *et al*., 2014). Functionally destabilizing mutations can be used in diagnostic panels (van Veen *et al*., 2018), as prognostic indicators of treatment response (Shieh *et al*., 2017), and even as targets for protein modulation as has been shown for CFTR in Cystic Fibrosis treatments (Pettit and Fellner, 2014). No single mutation guarantees tumorigenesis in breast cancer. However, if we can understand key mutations that cause protein dysfunction and interfere with cellular regulation and stability, we likely will be able to better diagnose and treat the disease.

## Acknowledgements

We thank Sophia Tan for helpful comments on the manuscript. This work was done as part of a M.S. in Bioinformatics degree at Brandeis University for one of the authors (M.A.C.).

